# The Lorenz’s side of Gene Set Enrichment Analysis: Similarity and divergences of the Gene Set Enrichment Analysis from the measure of wealth inequality

**DOI:** 10.1101/2024.11.23.624984

**Authors:** Stefano Maria Pagnotta

## Abstract

In the age of microarrays (about 20 years ago), the poor quality of expression levels prompted the development of methods for associating a set of significant genes with their biological meaning. A fundamental shift in methodology was the consideration of the entire gene profile, summarizing the differential level of genes measured in two conditions. A preliminary proposal was to mimic the two samples Kolmogorov-Smirnov test. Still, the winning idea was to introduce a system of weights in constructing the empirical distribution functions. Since 2005, the Gene Set Enrichment Analysis has emerged as the most well-known methodology from these premises.

This method is always referred to as based on a weighted Kolmogorov-Smirnov test. Sometimes, what is introduced as an innovation that improves a standard methodology leads to a well-known tool in a different science subject. While the accumulation of counts generates the empirical distribution function, the accumulation of weights, as defined in GSEA, leads to a function known as the Lorenz curve, introduced in 1905. Such a tool is a cornerstone in welfare studies to measure the concentration or equidistribution of richness in populations.

This paper reviews the essentials of the Lorenz curve and Gene Set Enrichment Analysis. It shows that the test statistic of the last is linked to the null hypothesis comparing two Lorenz curves. The new light of the enrichment procedure makes consistent analytical tools and conceptual formulation of the methodology.

## 1 Introduction

Gene-set enrichment analysis (GSEA) [Subramanian et al., 2005] is a widely known method that helps uncover the biological characteristics associated with a gene profile. This method is straightforward and requires specific components to produce results. On one side is a gene profile, essentially a ranked list of intensities, with each intensity linked to a gene. Usually, this gene profile is derived from comparing two sets of samples, such as tumor versus normal or treatment versus control. When the list is ordered, one end aggregates the genes with the highest differential intensity in the tumor group, while the other gathers the genes associated with the normal group. The second component is a set of genes, denoted as S, involved in a specific biological function or other characteristics. After obtaining the gene profile and the gene set, the crucial step is examining the gene profile with the gene set to determine whether the genes in the set are predominantly located on the treatment/tumor side or the control/normal side. This task is performed using a test statistic. Typically, a resampling algorithm is applied to compute the p-value, which determines the significance of the location of the gene set.

In this paper, we discuss GSEA test statistics. It comes as a generalization of a previous proposal from almost the same authors [Mootha et al., 2003]. Initially, they considered the Kolmogorov-Smirnov (KS) two-sample test statistics [Hollander et al., 2015, pp.190-194] to solve a test with *ℋ* _0_ : *F*_*in*_(*x*) = *F*_*out*_(*x*), ∀*x. F*_*in*_(*x*) is the population version of the distribution function (df) associated with the genes “in” the gene set; *F*_*out*_(*x*) is instead the df concerning the genes “out” the gene set [Debrabant, 2017]. When the null hypothesis is not rejected, the gene set thought as a unit gene [Wu and Smyth, 2012] cannot be associated with the treatment or control group. The implicit alternative hypothesis *ℋ* _1_ : ∃*x* s.t. *F*_*in*_(*x*) *≠ F*_*out*_(*x*) covers different cases that can lead to the rejection (see also [Barry et al., 2008]). That of interest for enrichment analysis is *F*_*in*_(*x*) *< F*_*out*_(*x*) that moves the original KS test in the field of first-order rule of stochastic dominance [Levy, 1992]. In this case, the gene set covers genes having the highest differential expression, leaving a significant association of the gene set with the treatment/tumor group. This latter shape of the alternative hypothesis, together with *H*_0_, is the system of hypotheses studied by Mann and Whitney [1947].

The improvement proposed in Subramanian et al. [2005] is the weighted KS-like test statistic. Given a set of observed values, *x*_1_, *x*_2_, …, *x*_*n*_, the empirical distribution function

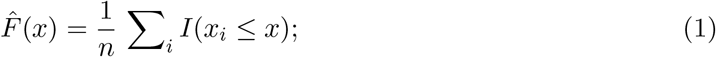

can be described as the percentage of the observed values not greater than *x*. The weighted KS-like test statistic 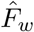 (*x*) is similar to edf, but the proportion of observed intensities not greater than *x* are computed instead of frequencies and apply to positive data only.

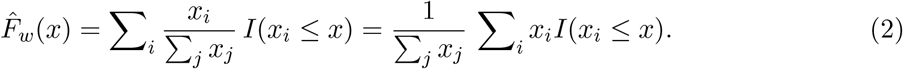

*I*(·) is the indicator function. 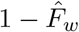 is a key element of the GSEA test statistic.

Lorenz [1905] has introduced 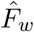 as a tool to measure a population’s wealth diffusion or concentration. Wealth can be measured by a positive statistical variable such as income. If the *x*_*i*_’s measure the families’ income, and we assume that *x*_1_ ≤ *x*_2_ ≤ · · · ≤ *x*_*n*_, then 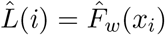 is the percentage of wealth owned by the 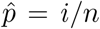 percentage of the poorest families. When the income is the same for all families, the 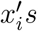*s* are constant, then 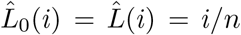. The sequence of segments connecting the points 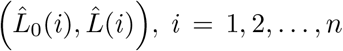, *i* = 1, 2, …, *n* is known as Lorenz curve [Gastwirth, 1971]. Figure 1 shows the Lorenz curve (the solid line) for 500 positive toy values. Two times the measure of the area (the shaded surface) between the Lorenz curve and the bisector is a concentration measure.

**Figure 1.**
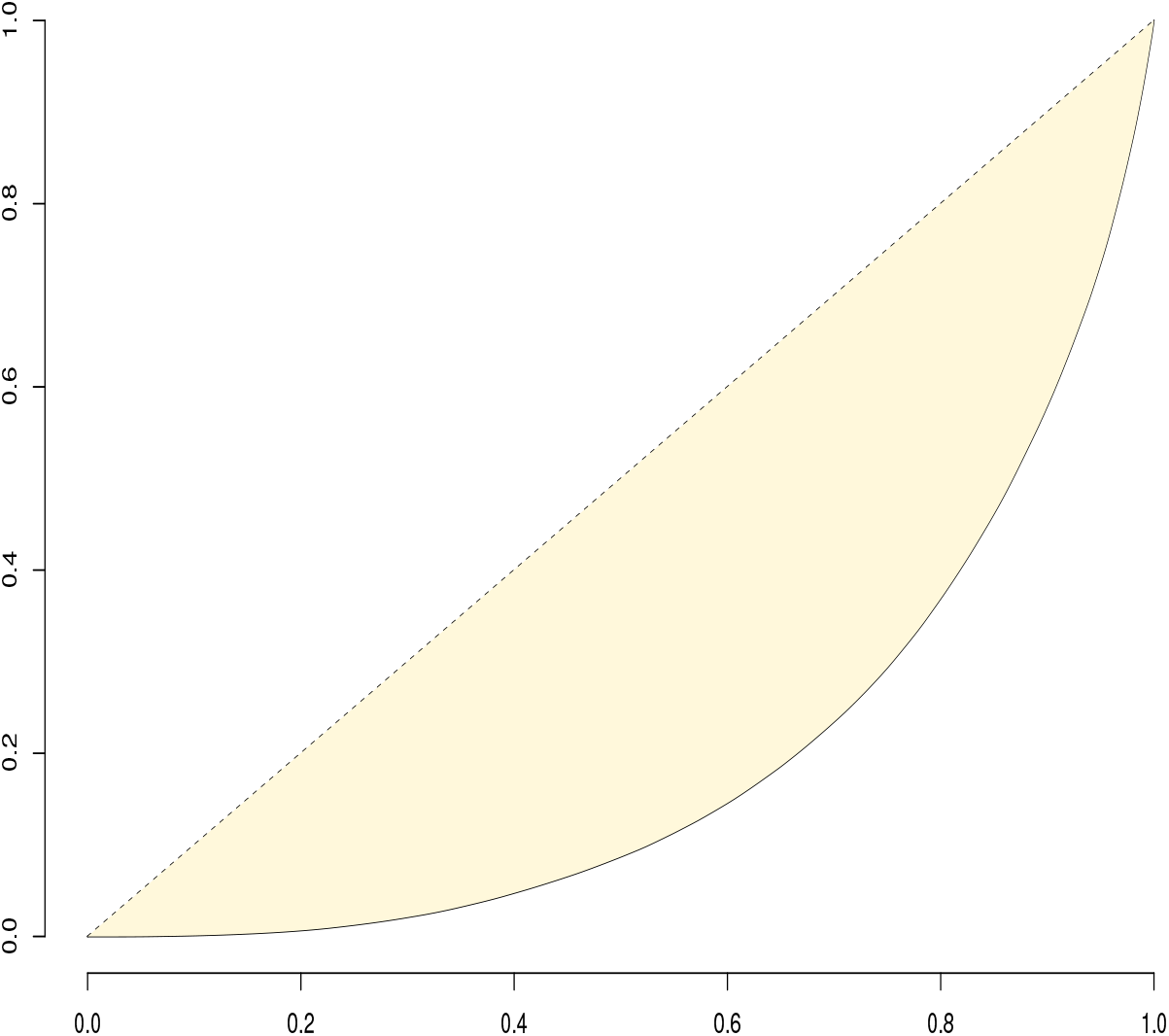
Lorenz curve (solid line) for 500 positive toy values. The shaded surface is the concentration area.

Since 1905, many studies in the economic and social fields have been carried out to move from a graphical display of the Lorenz curve to metrics. These fields are still under research attention [Sitthiyot and Holasut, 2021]. One of the most interesting results is the analytical link between Lorenz curve and *Gini’s concentration index* (GCI) [Giorgi and Gigliarano, 2017]

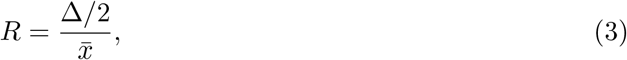

where

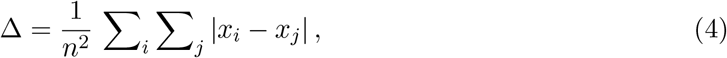

that is the *Gini’s Mean Difference* and falls in the class of the measures of variability [Yitzhaki, 2003]. Section 2 clarifies that the GCI is connected with the concentration area.

When the statistical units are the genes in a sample, and the wealth is identified with the differential expression level, the *rationale of grouped genes* (the gene-set) *analysis* section in Mootha et al. [2003] sounds so similar to a subject of concentration of expression levels in a few genes participating to a biological function. They observe that all the genes in the gene set are not expected to have “*fully concordant expression levels, but as long as an appreciable fraction of our gene set has this property, we can expect a benefit from the grouped gene approach*.” The adoption of (2) in Subramanian et al. [2005] is in the direction to amplify the effect of the differential expression level of some genes in the gene-set to the total of genes. This is not deliberately a question of measuring how much the total differential expression level of genes in a gene set is concentrated on them, but this is the point.

Before continuing, it is important to point out that values in the differential expression can be both positive and negative, but in the GSEA test statistic, they come as absolute values, so they are only positive.

In this note, we show that 1) Gene Set Enrichment Analysis is theoretically connected to the Lorenz curve, 2) the test statistic of GSEA is a fisherian tool to decide about the equality of two Lorenz curves.

The remainder of the article is organized as follows. In Section 2, we review the analytical results of the Lorenz curve. In Section 3, we outline the essentials of the GSEA methodology. Section 4 discusses the results of the previous two sections, and the conclusions are in Section 5.

## 2 The Lorenz curve

This section’s results are consistent with the notation from Gastwirth [1972].

Given a sequence of real positive values *x*_1_, *x*_2_, …, *x*_*n*_, being *x*_*i*_ ≤ *x*_*i*+1_, the empirical Lorenz curve (LC) is the graph of

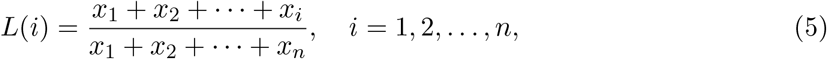

where the abscissa is *L*_0_(*i*) = *i/n. L*_0_ is the LC when *x*_1_ = *x*_2_ = · · · = *x*_*n*_ *>* 0. *L*(*i*) is the summation of the intensities *x*_1_, *x*_2_, …, *x*_*i*_ to the total amount 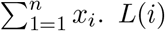 is always a value between 0 and 1, and *L*(*i*) ≤ *L*(*i* + 1), by construction. The graph is a convex curve. Figure 2a shows an example of the curve coming from toy values. Intuitively, if the *x*_*i*_’s are the total income of families in the population, a point on the Lorenz curve (LC) quantifies the proportion *L*(*i*) of the total income of the poorest 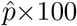 percent of the population, being 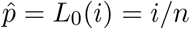.

**Figure 2.**
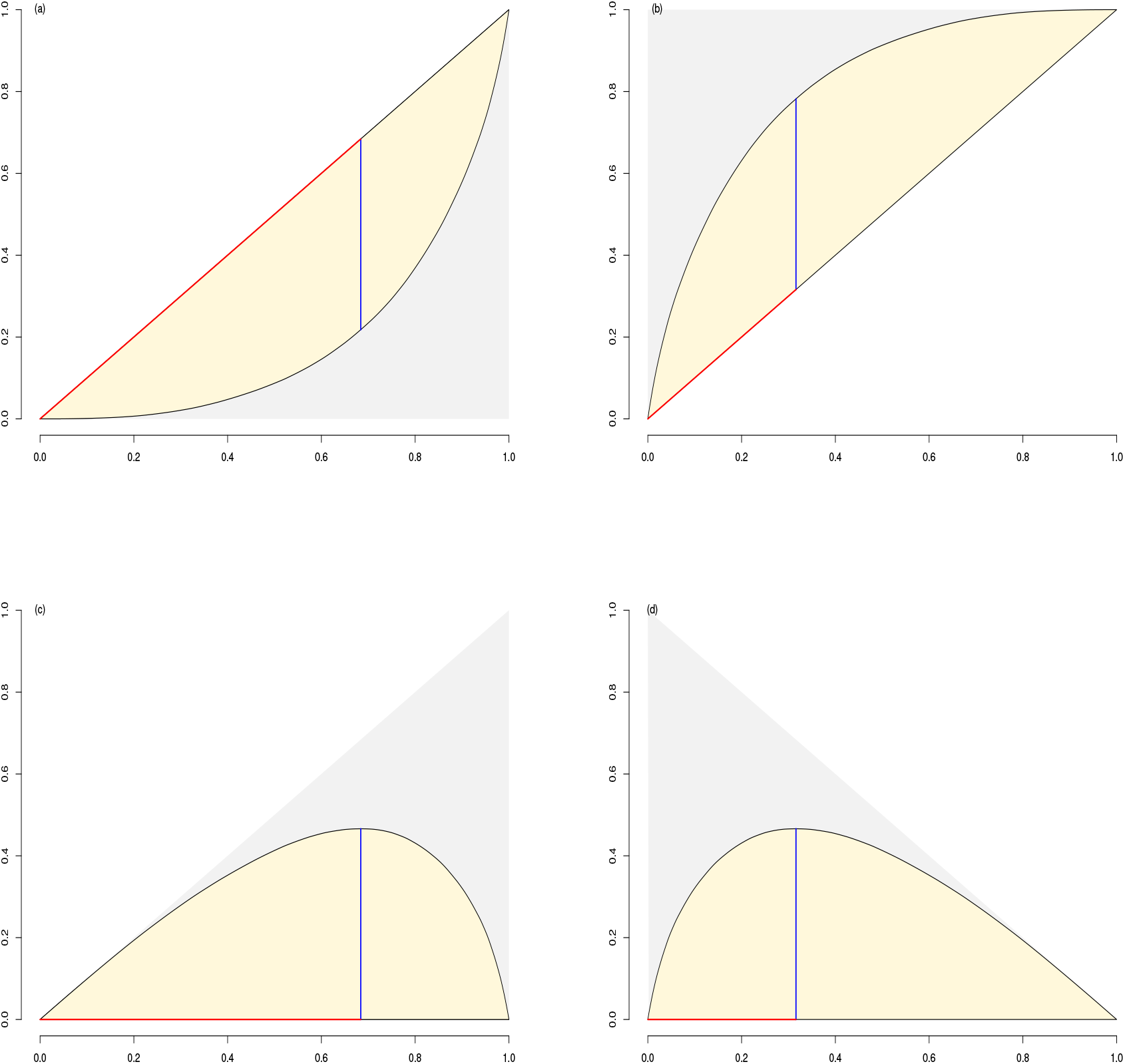
(a) Lorenz curve for 500 toy positive values; (b) Lorenz’s curve when the values are ordered from the largest to the smallest; (c) graph of discrepancy *L*_0_−*L*; (d) graph of discrepancy 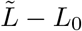 when the values are ordered from the largest to the smallest. In any graph, the shaded area is a measure of concentration, the right end of the red segment is the point of the maximum discrepancy, and the length of the blue segment is the maximum discrepancy.

If *F* (*x*) is the distribution function (with positive support) generating the values *x*_*i*_ from the random variable *X*, and *Q*(*p*) = *F* ^−1^(*p*) is the corresponding quantile function, LC admits [Gastwirth, 1971] a theoretical representation as

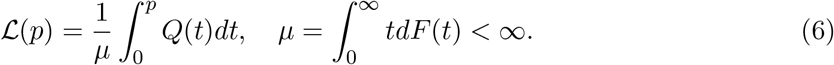

By applying the transformation *p* = *F* (*x*), the LC can be reparametrized with respect to *x* as follows

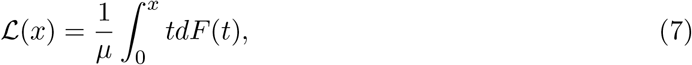

where *x* ranges in the support of *X*. This version of LC is the population counterpart of (2).

The Lorenz curve is connected to the measures of concentration. The *concentration area* is the area between *L*_0_ and *L* (shaded in Figure 2a, and the others). Its measure is

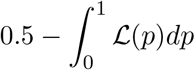

The ratio of the concentration area to the area of the lower triangle (grey-shaded)

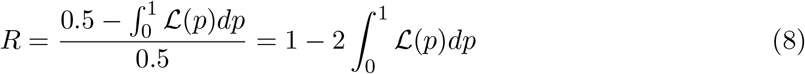

is the *Gini concentration index* [Kendall, 1948, prop.2.21], as proved by Gastwirth [1972] and clarified by Giorgi and Gigliarano [2017] and Yitzhaki [2003]. Equating (8) with the population version of (3) suggests that there is an analytical connection between LC and a measure of variability, actually the Gini’s mean difference.

Some other metrics can come from LC. Let

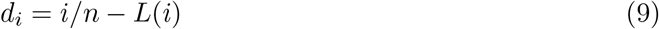

be the discrepancy of *L*_0_ from LC. Figure 2c shows the graph of the discrepancy. The percentile *p*^⋆^ = *i*^⋆^*/n* that maximizes *d*_*i*_, for *i*, is called *point of maximum discrepancy* (the right end of the red segment in Figure 2a); *i*^⋆^*/n* − *L*(*i*^⋆^) is instead the *maximum discrepancy* (the length of the blue segment in Figure 2a). The maximum discrepancy is itself another metric of concentration falling in the class of the *relative inequalities* [Gastwirth, 1972, *lemma 3]. In the theoretical setup, with basic calculus, it can be shown that the point p*^⋆^ of maximum discrepancy is *p*^⋆^ = *F* (*µ*) (see Lemma 2 in the appendix).

To match the results of the next section, we explore how the properties of LC change when the ordering of *x*_*i*_’s in (5) is reversed. Let 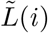(*i*) the version of (5) where *x*_1_ ≥ *x*_2_ ≥ · · · ≥ *x*_*n*_.

The graph of 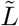 is concave (see Figure 2b). The measure of the area between 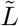 and *L*_0_ is the same as in the case of *L* and *L*_0_ (see Lemma 3 in the appendix; compare Figures 2a and 2b), then the Gini concentration index (GCI) is the same as well. If we set the discrepancy as

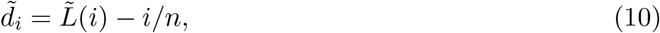

the maximum discrepancy of *L* and 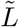 is the same (see Lemma 5), but the point of maximum discrepancy is 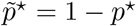 (see Lemma 4).

The theoretical equivalent of (6) is

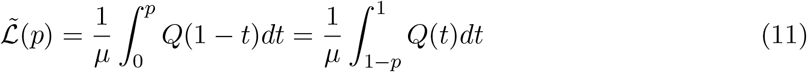

A trivial identity connects (6) and (11): appendix). 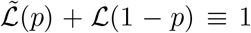 for all *p* (see Lemma 1 in the In Figures 3a and 3b, there are the plots of the 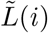 and of the corresponding discrepancy 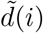 applied to real data. As an example, we considered the gene profile in Cerulo and Pagnotta [2022] and the OXIDATIVE PHOSPHORYLATION gene set in the Hallmark collection from MsigDB [Liberzon et al., 2011]. This gene list recaps the differential profiles of samples carrying the FGFR3-TACC3 gene-fusion positive (say, treatment group, with the corresponding gene at the top) versus the fusion negative (control group).

**Figure 3.**
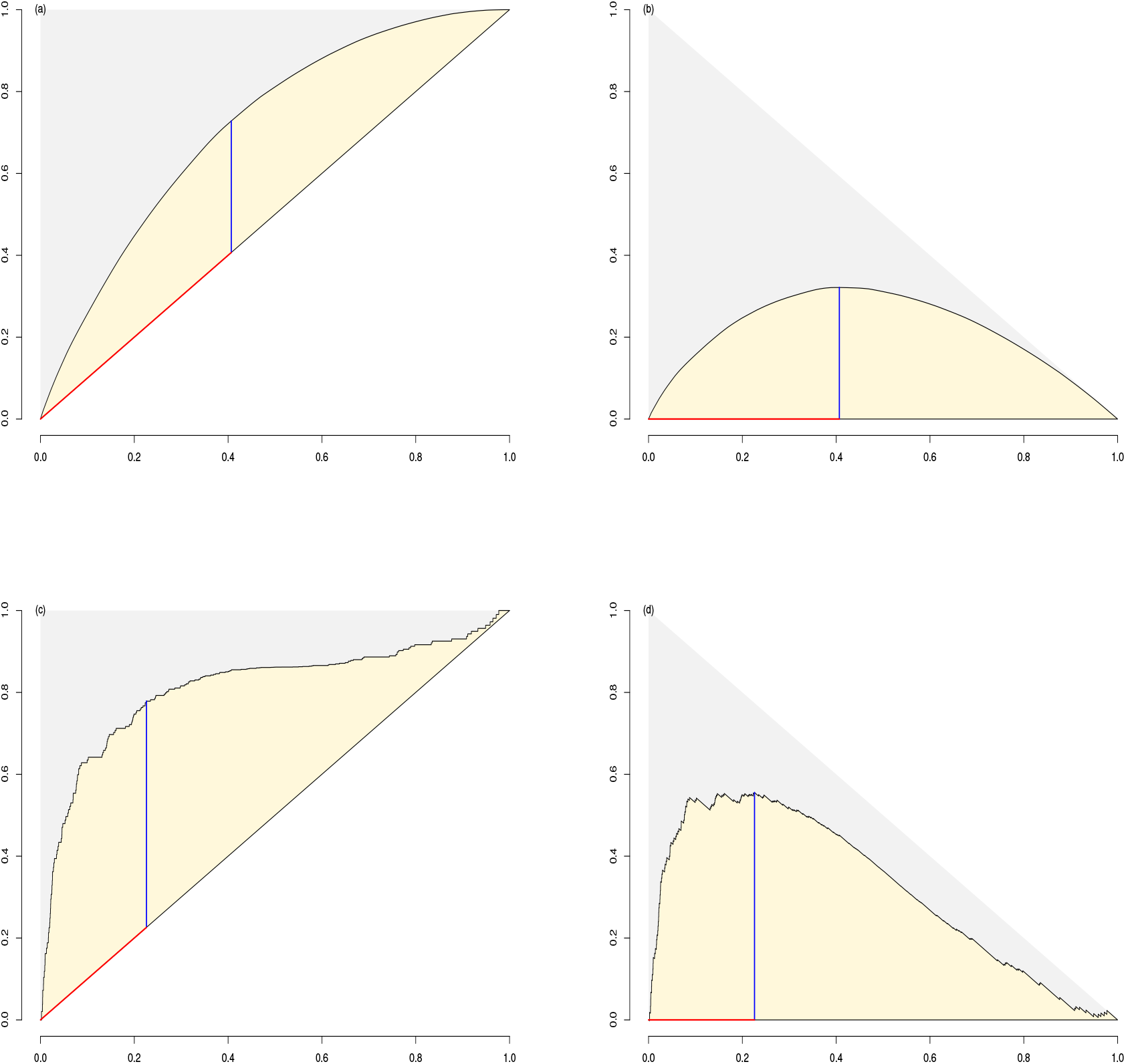
Graphs associated with a real ranked list of genes queried by the OXIDATIVE PHOSPHORYLATION gene-set in the Hallmark collection from MsigDB. (a) and (b) plots of 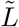 and the corresponding discrepancy 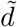; (c) is the plot of P_*hit*_, and (d) the plot of the running sum *RS*_*i*_ versus *i/N*. In (a) and (b), the right end of the red segment is the point of the maximum discrepancy, and the length of the blue segment is the maximum discrepancy. In (c) and (d), the red segment contains the leading edge subset of genes, and the length of the blue segment is the enrichment score.

In this section, we formally introduced the Lorenz curve and its most relevant results. We explored the impact of reversing the ordering of the values generating the LC and concluded that almost everything is preserved except the point of maximum discrepancy.

## 3 The Gene Set Enrichment Analysis statistic

We follow the original notation of Subramanian et al. [2005] where possible. Let *r*_1_, *r*_2_, …, *r*_*N*_ be the ordered values of the ranked list of genes, i.e. · · · ≥ *r*_*j*_ ≥ *r*_*j*+1_ ≥ · · ·, where *N* is the length of the list. Think of the *r*_*j*_’s as the differential gene expression levels coming from the comparison of two groups of samples. The genes with the highest *r*_*j*_ intensities (top genes) are in the list’s first locations. Given a gene-set *S*, the GSEA methodology needs the introduction of two empirical functions

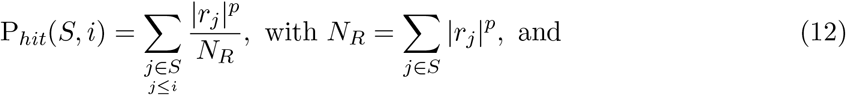

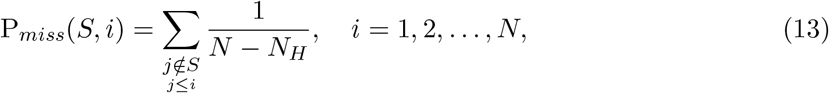

*N*_*H*_ is the number of genes in the gene set *S*. P_*hit*_(*S, i*) is the summation of the firsts *i* terms |*r*_*j*_|^*p*^, with *j* ≤ *i*, to the summation of all the |*r*_*j*_|^*p*^ inside the. Instead, P_*miss*_(*S, i*) is the fraction of the |*r*_*j*_|^*p*^’s outside the gene-set at the *i*^*th*^ location.

To simplify, let *x*_*i*_ = |*r*_*i*_|^*p*^, *x*_1_ ≥ *x*_2_, ≥ · · ·, ≥ *x*_*N*_, (12) can be reformulated as follows

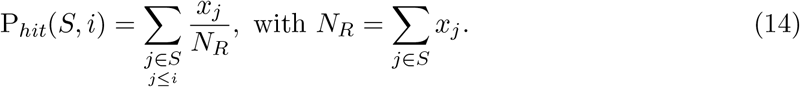

Let introduce the hit function *H*_*S*_(*x*) defined as

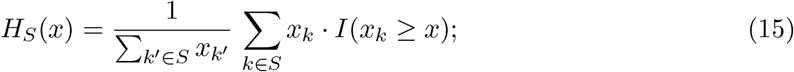

considering (2) it results 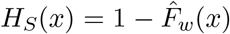 (see also Tamayo et al. [2016]). If *x*_*i*_ is a value of a gene in the gene set S, *H*_*S*_(*x*_*i*_) is the ratio of summation of the expression level of the genes in the gene set not lower than *x*_*i*_ to the summation of the expression levels of all gene in the gene-set. The value of *H*_*S*_(*x*) when *x*_*j*_ corresponds to a gene outside the gene set is *H*_*S*_(*x*_*i*_), where *x*_*i*_ is the closest value in the gene set such that *x*_*i*_ *> x*_*j*_. Finally, it follows that

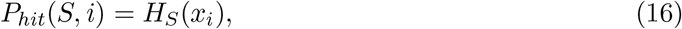

that is *P*_*hit*_(*S, i*) is *H*_*S*_ evaluated in the i^*th*^ value of the ordered sequence of *x*_*i*_’s. *H*_*S*_ is an increasing step function^1^, with jumps occurring when *x* is any of the values associated with the gene set.

When *H*_*S*_ is evaluated in the values associated with the genes in the gene set, then its values are the same as LC computed with the intensities, ordered from the largest to the smallest, that is *H*_*S*_(*x*_*i*_) = 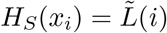(*i*) with *i* ∈ *S*. In this view, *H*_*S*_(*x*) extends the LC to the values corresponding to genes outside the gene set.

The population version of 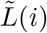 is 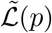 as defined in (11). Charmpi and Ycart [2015] reshape *P*_*hit*_(*S, i*) as LC, as shown in lemma (6). These authors talk of accumulated weights.

If we introduce

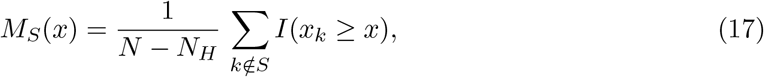

then *P*_*miss*_(*S, i*) is *M*_*S*_(*x*) evaluated at the observed value *x*_*i*_:

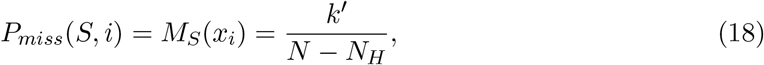

where *k*^*′*^ counts the number of *x*_*k*_ ≥ *x*_*i*_’s, *k* ∈*/ S*. When *M*_*S*_(*x*) is restricted to the values outside the set, then 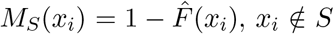, where 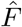 is the edf associated with the values of the genes outside the gene set. Being the 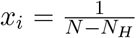 for all *i* ∈*/ S*, 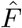 can be seen as the LC in the case of equidistribution, then 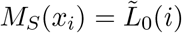.

From *P*_*hit*_ and *P*_*miss*_, a statistic is computed for each value in the gene profile:

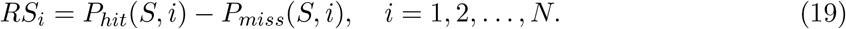

*RS*_*i*_, called running-sum, defines t he s tandard g raphical r epresentation o f G SEA a s t he plot of *RS*_*i*_ against *i*. In Figure 3d, we instead drew *RS*_*i*_ against *i/N*, which is a scaling of *i* to its maximum. Data are the same behind Figure 3b: a real gene profile interrogated by the OXIDATIVE PHOSPHORYLATION gene set. The length of the blue segment in Figure 3d is the metric of GSEA [Subramanian et al., 2005] called Enrichment Score (ES) and defined as the maximum deviation from zero of *P*_*hit*_ − *P*_*miss*_. The red segment contains the *leading edge subset of genes*, i.e., those genes in the gene set responsible for pushing the running sum to its maximum [Fleming and Miller, 2016, Tan et al., 2016].

The enrichment score is the test statistic to verify the significance o f *t he d egree t o w hich a set S is over-represented at the extremes (top or bottom) of the entire ranked list* as stated in Subramanian et al. [2005].

The running-sum can also be obtained as the evaluation of *H*_*S*_ (*x*) − *M*_*S*_ (*x*) at *x*_*i*_, that is

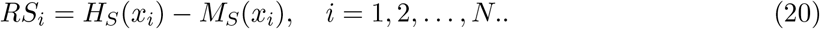

In this way, the maximum deviation from zero of *RS*_*i*_ follows the computation of the test statistic for the two samples of the Kolmogorov-Smirnov test, for example, in 5.4 (pp.192-194) of [Hollander et al., 2015].

## 4 Discussion

The seminal paper of Mootha et al. [2003] introduced a new approach to uncovering the biological features of two groups of samples. Previously, methods like the ones involving the hypergeometric distribution required only some genes to participate in this aim (see Boyle et al. [2004] for an example). The innovation was to involve all genes in the process instead of considering only those showing a significant differential expression and comparing them with a set of the co-regulated genes committed to some biological function. Given such a collection of genes, the gene set, they designed a test statistic connected with the Kolmogorov-Smirnov test. Referring to this method, they implicitly suggested that the aim is to compare the distribution functions associated with the gene level expression of those genes inside (consider *F*_*in*_) the gene set and the levels of genes outside (*F*_*out*_).

Exactly, the KS test null hypothesis requires that *F*_*in*_(*x*) = *F*_*out*_(*x*) for all *x*; the alternative hypothesis is instead that *F*_*in*_(*x*) ≠ *F*_*out*_(*x*) for at least one *x*. In the broad range of cases such that the alternative hypothesis is fulfilled, there is the chance *F*_*in*_(*x*) *< F*_*out*_(*x*) for all *x*. In this specific alternative, we find the answer to the location problem, as stated in Subramanian et al. [2005] when they say *The goal of GSEA is to determine whether members of a gene set S tend to occur toward the top (or bottom) of the list L, in which case the gene set is correlated with the phenotype class distinction*. If *F* (*x*) is a distribution function, its inflection point is the mode. If the KS null hypothesis is rejected towards *F*_*in*_(*x*) *< F*_*out*_(*x*), it means that the mode of the genes in the gene set is higher than that of the genes outside. The gene set is then associated with the group generating positive differential expression. The rejection of the KS null hypothesis is not necessarily connected to a change in the location of the two distributions [Barry et al., 2008].

The improvement of the original method is the weighted KS test. Essentially, in Subramanian et al. [2005], the test statistic is no longer based on empirical distribution functions, which are the accumulation of counts, but they consider a weighted accumulation of counts. The weights are a transformation of gene expression levels. As shown before, the weighted accumulation of counts results in an accumulation of intensities. The accumulation of intensities generates a Lorenz curve. The use of the Lorenz curves implies that the null hypothesis would now be

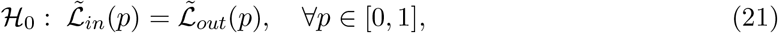

where 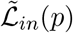 is the LC associated with the gene in the gene set, and 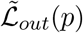 is the LC associated with the genes outside the gene set having the intensities all set to the same value. The shape of this hypothesis is also supported by the test statistic involving the maximum deviation of the two functions *H*_*S*_(*x*) and *M*_*S*_(*x*) that are both extensions of Lorenz curves.

The null hypothesis means there is no gene expression level concentration in the genes of the gene set. In different terms, it can be said that expression levels of genes inside the gene set are almost the same value. Then, the test is about variability and not location. Such an interpretation is supported by

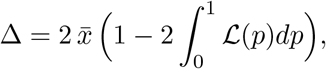

coming from (3), (4), (8), and lemma (3), where the LC is analytically connected to the Gini’s mean difference, that is a metric of scale.

The test statistic is the maximum deviation from zero of the running sum (19, 20). Such a statistic recalls the test statistic of the two samples KS test for comparing two populations. With a computational approach, the distribution of the test statistic is obtained under the null hypothesis, and the *p*-value is evaluated. Instead, inference about LC follows other ways [Beach and Kaliski, 1986].

## 5 Conclusion

We have examined the GSEA test statistic to emphasize its essential nature. The test statistic is the maximum deviation from zero of the running sum, as formalized in (19, 20). We have established a population version of *P*_*hit*_ and *P*_*miss*_ to obtain *H*_*S*_ and *M*_*S*_, which are Lorenz curves. The test statistic is the maximum deviation from zero of two Lorenz curves. The null hypothesis for this statistic is the equality of two Lorenz curves. This proposal is consistent with other contexts of hypothesis testing. For instance, the standardized difference of two sample means is the test statistic for the null hypothesis of equality of two population means. Also, the claim of mimicking the KS test presupposes the equality of two population distribution functions.

The GSEA methodology is deeply grounded in standard tools for measuring wealth inequality. The concept of concentration, as depicted in the Lorenz curves, aligns with the original idea of the GSEA authors, who expect to identify at least a few genes in the gene set with a higher differential level. Although the theoretical essence of GSEA revolves around variability, the test statistic allocates the gene set close to treatment or control groups of samples, adding a layer of complexity to the methodology.

## Acknowledgments

Stefano Maria Pagnotta is financially supported by PRIN 2022 PNRR grant n.P202228YH4 (CUP F53D23008680001).

## Appendix: some lemma. The shared setup of the following lemma is

1. *F* (*x*) is a distribution function with support *x* ≥ 0 and expected value *µ <* ∞
2. *Q*(*p*) = *F* ^−1^(*p*) is the quantile function associated with *F* (*x*)
3. the Lorenz curve is

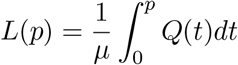
4. the Lorenz curve representing the ordering of values from the largest to the smallest is

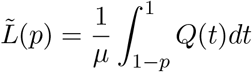

### Lemma 1.

*Under points 1-4, it results*

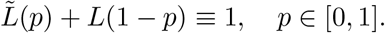

*Proof*.

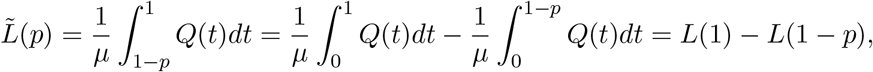

being *L*(1) = 1, it comes the thesis.

### Lemma 2.

*Under points 1-3, the point of maximum discrepancy is p*^⋆^ = *F* (*µ*).

*Proof*. Let

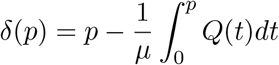

be the discrepancy of *L*_0_ from *L*, it follows that

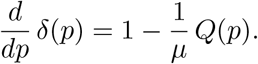

By applying the necessary condition for extremum, it results in the equation

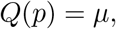

whose solution is

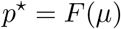

that is *p*^⋆^ a point of minimum, being *δ*(*p*) a concave function by construction.

### Lemma 3.

*Under points 1-4, it results that the measure of the area between L*_0_ *and L*

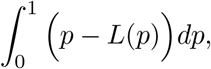

*and the measure of the area between* 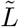 *and L*_0_

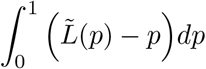

*are the same*.

*Proof*. With Lemma 1, it follows

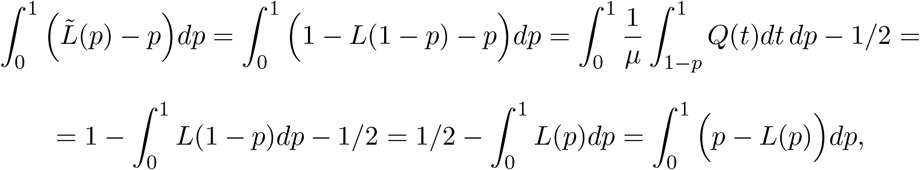

Being

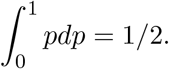

### Lemma 4.

*Under points 1-4, it results that the point of maximum discrepancy associated with*

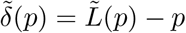

*is* 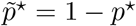, *being p*^⋆^ = *F* (*µ*).

*Proof*. With Lemma 1, it follows

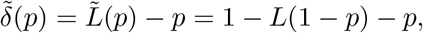

Then

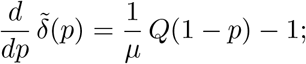

by applying the necessary condition for extremum it results in the equation

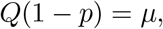

whose solution is

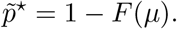

From Lemma 2, it follows

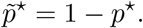

### Lemma 5.

*Under points 1-4, it results that the maximum discrepancy associated with*

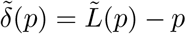

*is the same of*

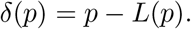

*Proof*. 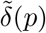 is maximum in 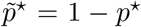, while *δ*(*p*) is maximum in *p*^⋆^. By using the Lemma 1 and 4, it results

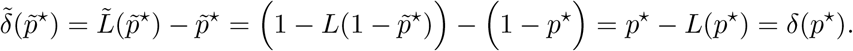

### Lemma 6.

*The function*

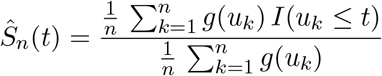

*of [Charmpi and Ycart, 2015, (equation 1)] is a Lorenz curve*.

*Proof*. From Bullock [1988], by using the Riemann-Stieltjes integral, it comes to the identity

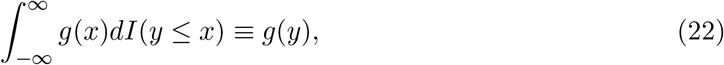

where *I*(·) is the indicator function, i.e., its value is 1 when the argument is true, 0 otherwise; *g*(*x*) is a continuous function. With the plug-in principle, the Lorenz curve 7

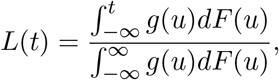

where *g*(*u*) = *u*, can be approximated as

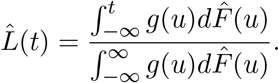

With the identity 22, it follows that

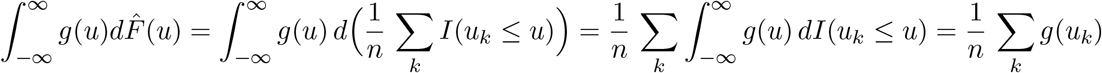

And

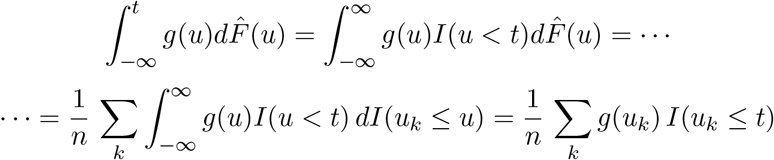

Then

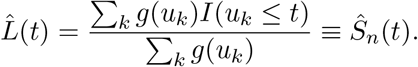

## Bibliography

W. T. Barry, A. B. Nobel, and F. A. Wright. A statistical framework for testing functional categories in microarray data. The Annals of Applied Statistics, 2(1):286 – 315, 2008. doi: 10.1214/07-AOAS146. URL https://doi.org/10.1214/07-AOAS146.

C. M. Beach and S. F. Kaliski. Lorenz curve inference with sample weights: An application to the distribution of unemployment experience. Journal of the Royal Statistical Society. Series C (Applied Statistics), 35(1):38–45, 1986. ISSN 00359254, 14679876. URL http://www.jstor.org/stable/2347862.

E. I. Boyle, S. Weng, J. Gollub, H. Jin, D. Botstein, J. M. Cherry, and G. Sherlock. GO::TermFinder—open source software for accessing Gene Ontology information and finding significantly enriched Gene Ontology terms associated with a list of genes. Bioinformatics, 20(18):3710–3715, 08 2004. ISSN 1367-4803. doi: 10.1093/bioinformatics/bth456. URL https://doi.org/10.1093/bioinformatics/bth456.

G. L. Bullock. A geometric interpretation of the riemann-stieltjes integral. The American Mathematical Monthly, 95(5):448–455, 1988. ISSN 00029890, 19300972. URL http://www.jstor.org/stable/2322483.

L. Cerulo and S. M. Pagnotta. massivegst: A mann-whitney-wilcoxon gene-set test tool that gives meaning to gene-set enrichment analysis. Entropy, 24(5), 2022. ISSN 1099-4300. doi: 10.3390/e24050739. URL https://doi.org/10.3390/e24050739.

K. Charmpi and B. Ycart. Weighted kolmogorov smirnov testing: an alternative for gene set enrichment analysis. Statistical Applications in Genetics and Molecular Biology, 14 (3):279–293, 2015. doi: doi:10.1515/sagmb-2014-0077. URL https://doi.org/10.1515/sagmb-2014-0077.

B. Debrabant. The null hypothesis of GSEA, and a novel statistical model for competitive gene set analysis. Bioinformatics, 33(9):1271–1277, Jan. 2017.

D. S. Fleming and L. C. Miller. Leading edge analysis of transcriptomic changes during pseudorabies virus infection. Genomics Data, 10:104–106, 2016. ISSN 2213-5960. doi: 10.1016/j.gdata.2016.09.014. URL https://www.sciencedirect.com/science/article/pii/S2213596016301295.

J. L. Gastwirth. A general definition of the lorenz curve. Econometrica, 39(6):1037–1039, 1971. ISSN 00129682, 14680262. URL http://www.jstor.org/stable/1909675.

J. L. Gastwirth. The estimation of the lorenz curve and gini index. The Review of Economics and Statistics, 54(3):306–316, 1972. ISSN 00346535, 15309142. URL http://www.jstor.org/stable/1937992.

G. M. Giorgi and C. Gigliarano. The gini concentration index: A review of the inference literature. Journal of Economic Surveys, 31(4):1130–1148, 2017. doi: 10.1111/joes.12185. URL https://onlinelibrary.wiley.com/doi/abs/10.1111/joes.12185.

M. Hollander, D. A. Wolfe, and E. Chicken. Nonparametric Statistical Methods. John Wiley & Sons, Ltd, 2015. ISBN 9781119196037. doi: 10.1002/9781119196037.ch4. URL https://onlinelibrary.wiley.com/doi/abs/10.1002/9781119196037.ch4.

M. G. Kendall. The Advanced Theory of Statistics, volume 1. C. Griffin, London, 1948.

H. Levy. Stochastic dominance and expected utility: Survey and analysis. Management Science, 38(4):555–593, 1992. ISSN 00251909, 15265501. URL http://www.jstor.org/stable/2632436.

A. Liberzon, A. Subramanian, R. Pinchback, H. Thorvaldsdóttir, P. Tamayo, and J.P. Mesirov. Molecular signatures database (MSigDB) 3.0. Bioinformatics, 27(12):1739–1740, 05 2011. ISSN 1367-4803. doi: 10.1093/bioinformatics/btr260. URL https://doi.org/10.1093/bioinformatics/btr260.

M. O. Lorenz. Methods of measuring the concentration of wealth. Publications of the American Statistical Association, 9(70):209–219, 1905. doi: 10.1080/15225437.1905.10503443. URL http://www.tandfonline.com/doi/abs/10.1080/15225437.1905.10503443.

H. Mann and D. Whitney. On a test of whether one of two random variables is stochastically larger than the other. Ann. Math. Statist., 18(1):50–60, 03 1947. doi: 10.1214/aoms/1177730491.

V. K. Mootha, C. M. Lindgren, K.-F. Eriksson, A. Subramanian, S. Sihag, J. Lehar, P. Puigserver, E. Carlsson, M. Ridderstråle, E. Laurila, N. Houstis, M. J. Daly, N. Patterson, J. P. Mesirov, T. R. Golub, P. Tamayo, B. Spiegelman, E. S. Lander, J. N. Hirschhorn, D. Altshuler, and L. C. Groop. Pgc-1α-responsive genes involved in oxidative phosphorylation are coordinately downregulated in human diabetes. Nature Genetics, 34(3):267–273, Jul 2003. ISSN 1546-1718. doi: 10.1038/ng1180. URL https://doi.org/10.1038/ng1180.

T. Sitthiyot and K. Holasut. A simple method for estimating the lorenz curve. Humanities and Social Sciences Communications, 8(1):268, Nov 2021. ISSN 2662-9992. doi: 10.1057/s41599-021-00948-x. URL https://doi.org/10.1057/s41599-021-00948-x.

A. Subramanian, P. Tamayo, V. K. Mootha, S. Mukherjee, B. L. Ebert, M. A. Gillette, A. Paulovich, S. L. Pomeroy, T. R. Golub, E. S. Lander, and J. P. Mesirov. Gene set enrichment analysis: A knowledge-based approach for interpreting genome-wide expression profiles. Proceedings of the National Academy of Sciences, 102(43):15545–15550, 2005. doi: 10.1073/pnas.0506580102.

P. Tamayo, G. Steinhardt, A. Liberzon, and J. P. Mesirov. The limitations of simple gene set enrichment analysis assuming gene independence. Statistical Methods in Medical Research, 25(1):472–487, 2016. doi: 10.1177/0962280212460441. URL https://doi.org/10.1177/0962280212460441.

Y. Tan, J. Godec, F. Wu, P. Tamayo, J. P. Mesirov, and W. N. Haining. A method for downstream analysis of gene set enrichment results facilitates the biological interpretation of vaccine efficacy studies. bioRxiv, 2016. doi: 10.1101/043158. URL https://www.biorxiv.org/content/early/2016/04/11/043158.

D. Wu and G. K. Smyth. Camera: a competitive gene set test accounting for inter-gene correlation. Nucleic Acids Research, 40(17):e133–e133, 05 2012. ISSN 0305-1048. doi: 10.1093/nar/gks461. URL https://doi.org/10.1093/nar/gks461.

S. Yitzhaki. Gini’s mean difference: a superior measure of variability for non-normal distributions. METRON - International Journal of Statistics, LXI(2):285–316, 2003. URL https://ideas.repec.org/a/mtn/ancoec/030208.html.

